# Brain Aging Among Individuals with Classical Trigeminal Neuralgia

**DOI:** 10.1101/2024.11.13.623489

**Authors:** Yenisel Cruz-Almeida, Pedro A. Valdes-Hernandez, Yun Liang, Mingzhou Ding, John K. Neubert

## Abstract

Trigeminal neuralgia (TN) is a complex orofacial neuropathic pain condition with limited understanding of underlying mechanisms and therapeutic options. Emerging evidence suggests the involvement of the brain in persons with TN including widespread brain changes when employing a widely used brain aging biomarker that estimates a predicted brain age difference or brain age gap. The aim of the present cross-sectional study was to assess the predicted brain age difference (brain-PAD) or brain age gap across two discrete TN subtypes (classical TN, and secondary/idiopathic TN) in comparison with age-and sex-matched pain-free controls and its association with several clinical and psychological characteristics. Thirty-four individuals diagnosed with Classical TN, 17 diagnosed with secondary/idiopathic TN were age- and sex-matched to pain-free controls (n=54). All participants underwent a T1 brain MRI and completed clinical and psychological measures. There were significant differences in brain-PAD among TN subtypes (ANCOVA p = 0.0078, effect size f^2^ = 0.28^2^), with individuals diagnosed with Classical TN having a brain-PAD significantly greater than the controls by 3.87 years (p = 0.01, Bonferroni-corrected). There were no significant brain-PAD differences between secondary/idiopathic TN and pain-free controls. Brain-PAD had a significant positive association with both pain catastrophizing (p = 0.032) and pain-related anxiety (p = 0.041), but no significant association with disease duration (p = 0.519) or usual pain intensity (p = 0.443). We report here accelerated brain aging processes in patients with classical TN, but not in persons diagnosed with secondary/idiopathic TN. Our study replicates previous findings and adds to the literature that accelerated brain aging may not occur across all TN subtypes. Given the increased use of MRI for TN diagnostics, combined with our own recent work deriving our brain aging biomarker from clinical-grade scans, future studies within clinical settings are feasible and needed to understand this debilitating condition.

## 1. Introduction

Trigeminal neuralgia (TN), previously known as tic douloureux, is a complex neuropathic pain condition characterized by recurrent brief episodes of electric, lancinating, shock-like pains affecting the structures innervated by the fifth cranial nerve. The prevalence of TN increases with age and women are disproportionately affected. The International Headache Society (IHS) and International Association for the Study of Pain (IASP) established a new classification system for TN, which includes subtypes of classical TN, secondary TN, and/or idiopathic TN (Olesen 2018). This classification scheme aims to incorporate the current understanding of the pathophysiology of the disease with clinical presentation to better aid clinicians in effectively diagnosing TN subtypes and thereby guiding treatments based on the diagnosis. Classical TN is generally driven by compression of the nerve root and typically responds to surgical intervention (e.g., microvascular decompression (MVD)), while secondary/idiopathic TN has no clear cause in most cases and may be more refractory to surgery. Given the lack of therapeutic options, research is urgently needed to understand the underlying neurobiology of the various TN subtypes to identify novel targets and improve treatments for this complex, debilitating condition.

Emerging evidence supports changes in brain structure and function of specific cortical and subcortical regions in TN patients compared to controls (Zhang et al. 2020). Further, individuals with TN also appear to have widespread brain changes when employing a widely used brain aging biomarker (Hung et al. 2022). However, the latter study did not differentiate between TN subtypes, nor examined the association between the brain age biomarker with measures of pain and psychosocial function. It is not currently clear whether accelerated brain aging processes occur across all TN diagnoses or only a specific subtype and the relationship with various clinical characteristics. Given the need to understand the pathophysiology of TN to guide treatment, the aim of the present study was to assess the predicted brain age difference or brain age gap across two discrete TN subtypes (classical TN, and secondary/idiopathic TN) in comparison with age-and sex-matched pain-free controls. We also examined whether the predicted brain age difference or brain age gap was associated with several clinical and psychological characteristics. We hypothesized that individuals with TN would have an older appearing brain compared to pain-free controls and that worse clinical and psychological characteristics would be associated with an older brain.

## 2. Materials and Methods

### 2.1. Participants

The present investigation was performed at the University of Florida (UF) Health Sciences Center (UFHealth) and the Evelyn F. and William L. McKnight Brain Institute (MBI) at the University of Florida. The study was carried out in accordance with the Declaration of Helsinki and the study conforms to the STROBE Guidelines. Screening was done either in person or via the telephone for all participants. All participants provided verbal and written informed consent during their visit to UFHealth. A trained study coordinator read a standard script explaining all the study procedures to all subjects.

#### 2.1.1. Individuals with Trigeminal Neuralgia

Participants with trigeminal neuralgia (TN) were recruited through the clinical care population at the University of Florida Health System from referrals provided by the Facial Pain Research Foundation. The study was approved by the WCG Institutional Review Board (IRB). Participants with TN completed the Oregon Health Science University (OHSU) Trigeminal Neuralgia – Diagnostic Questionnaire. A trained clinical fellow completed a focused medical history, a trigeminal cranial nerve exam, and a physical examination. Eligible subjects completed a study packet followed by an MRI scan. The study packet contained the following self-reported measures:

a. **Usual Pain Intensity:** Participants were asked to rate their daily pain experienced during the past month. Subjects indicated on a 100mm visual analog scale (VAS) anchored on the left with “no pain sensation” and on the right with “most intense pain sensation imaginable”.
b. **Pain Anxiety Symptom Scale (PASS):** Participants were asked to rate the frequency of responses to pain on a 0 (never) to 5 (always) rating scale. The PASS contains 4 subscales characterizing different responses to pain: cognitive, escape/avoidance, fear, and physiological anxiety (McCracken et al. 1992).
c. **Beck Depression Inventory-II (BDI-II):** The BDI-II measures the severity of depressive symptoms. The inventory is composed of 21 items relating to symptoms such as hopelessness and irritability, cognitions such as guilt or feelings of being punished, as well as physical symptoms such as fatigue, weight loss, and lack of interest in sex (Dozois et al. 1998).
d. **Pain Catastrophizing Scale (PCS):** The PCS consists of 13 statements containing a number of thoughts and feelings one may experience when having pain. Each item is scored on a 5-point scale (Sullivan et al. 1995).

#### 2.1.2. Pain-Free Controls

Healthy subjects who participated in previous studies at the UF PAIN laboratory and had previously consented for the reuse of their data for future studies were eligible to be included as controls. Control participants were defined as: 1) participants without reporting chronic pain on most days within the past 6 months using the Graded Chronic Pain Scale, and 2) had also MRIs obtained using the same scanner and MRI protocol as the TN participants. To match a TN participant to a control, we selected a random participant in our TN sample and matched them with the closest same sex participant in age from our control dataset. This procedure was then repeated for the rest of the participants excluding the ones already selected (i.e., without replacement). To ensure enough matching controls across all age ranges, the procedure prioritized those age groups where our control dataset had fewer participants. Age ranges with a larger participant pool were addressed last. This approach was adopted to prevent a potential shortage of matching controls in the less-represented age groups. Previous studies were approved by UF IRB01.

### 2.2. MRI data for TN and Controls

All MRI data was collected at the UF MBI using a 3-Tesla Achieva Phillips (Best, the Netherlands) scanner using a 32-channel radio-frequency coil. A high resolution, T1-weighted (T1w) turbo field echo anatomical image was collected with TR = 7.0 ms, TE = 3.2 ms, 176 slices acquired in a sagittal orientation, flip angle = 8 degrees, resolution = 1 mm^3^. Head movement was minimized via cushions positioned inside the head coil. Vital signs (blood pressure, temperature, and pulse) were also recorded prior to scanning.

### 2.3. Brain age estimations

The predicted brain age difference (brain-PAD) sometimes also called brain age gap (BAG) is the difference between a predicted brain age from an MRI and the person’s chronological age, and much research suggests its potential clinical value. Its utility to compare the degree of brain age acceleration among groups is straightforward. However, interpretability of absolute values, i.e., mean brain-PAD within a group, is obscured by the so-called “regression dilution” bias, which overestimates younger ages and underestimates older ages. This precludes us to understand which groups are actually experiencing accelerated brain aging and thus driving the group differences in brain-PAD. To report the clinically relevant unbiased brain-PADs, we recently advocated (Valdes-Hernandez, Laffitte Nodarse, Peraza, et al. 2023a; Valdes-Hernandez, Laffitte Nodarse, Cole, et al. 2023b) for the use of the correction of the predicted brain ages proposed by (Cole et al. 2018)

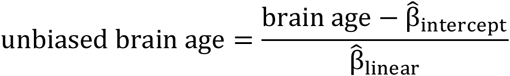

In this equation, 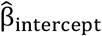 and 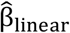 are the coefficients of the linear fit brain age∼chronological age.

### 2.4. Data and Statistical Analysis

The present study was powered based on our research (Cruz-Almeida et al. 2019; Valdes-Hernandez, Laffitte Nodarse, Johnson, et al. 2023) and others (Hung et al. 2022) where a sample size of 54 will provide over 90% power to detect a difference between two groups with two covariates at an alpha of 0.05. Missing data was treated using listwise elimination. To ensure normality, for every model (which entailed a specific subset of the whole dataset), we applied a rank-based inverse normal transformation to the dependent variable (brain-PAD) using the ‘Blom’ method with parameter c= 3/8 (Downton and Blom 1961). After fitting the models, we applied the Shapiro-Wilk test of composite normality (with unspecified mean and variance) on the residuals (for Platykurtic distributions; while the Shapiro-Francia test was used for Leptokurtic distributions) to test whether the normality assumption required for linear models was fulfilled (Shapiro and Wilk 1965). All analyses were re-run after removing those measurements deemed outliers, based on their Cook’s distance being 3 times higher than their sample average (NETER 1990).

To test whether there was a significant difference between groups in brain-PAD, we used Analysis of Covariance (ANCOVA) controlling for chronological age and sex. Thus, we tested the significance of the effect of *group* in the linear model *brain-PAD ∼ group + sex + age*, where *sex* took the values ‘male’ and ‘female’, and *age* is the chronological age. To be consistent with the study by Hung and colleagues (Hung et al. 2022), we first performed the analysis with the nominal categorical variable *group* with levels CONTROLS and TN. We performed a second analysis including group with levels CONTROLS, Classical TN and Secondary/Idiopathic TN. Effects sizes were reported using the Cohen’s f^2^, which measures the relative variance explained by the effect when added to the regression model (0.1^2^ ≤ f^2^ < 0.25^2^ for small effects, 0.25^2^ ≤ f^2^ < 0.4^2^ for medium effects and f^2^ > 0.4^2^ for large effects. For pairwise comparisons, we additionally reported the difference in marginal means, namely ΔPAD, and its Standard Error (SE). Statistical significance was set to α = 0.05, with p-values corrected for multiple comparisons using a Bonferroni correction. Finally, to examine whether brain-PAD was associated with clinical and psychological symptoms, we employed partial correlations corrected for multiple comparisons using the Benjamini-Hochberg procedure (*q*<0.05) (Benjamini and Hochberg 1995). Given there were significant differences between TN groups in BDI-II scores, BDI-II was included as a covariate in this analysis.

## 3. Results

### 3.1. Sample characteristics

The total sample is characterized in **Table 1**. In total, 55 TN participants gave written informed consent (69% female, mean age ± standard deviation (SD) = 53.9 ± 14.9). One participant did not meet diagnosis criteria (n = 1) for either classical TN or secondary/idiopathic TN and was excluded from the study. The data from the remaining 54 TN patients, and the 54 sex- and age-matched controls were analyzed and are reported here. Specifically, 34 individuals were diagnosed with Classical TN and 17 were diagnosed with secondary/idiopathic TN. As expected, chronological age was highly correlated with the predicted brain age across all participants (**Figure 1** and **Table 2)**.

**Table 1.** Characteristics of the sample (n = 108).

|  |  | Controls | Classical TN | Secondary/Idiopathic<br>TN | Statistics |
| --- | --- | --- | --- | --- | --- |
| Sex: | Females | 38 | 23 | 15 | $\chi^2 = 3.80$ |
| | Males | 16 | 14 | 2 | $p = 0.150$ |
| Age (years) | | 54.2<br>(17.4) | 57.7<br>(13.6) | 47.64<br>(14.2) | $F_{2,107} = 2.37$<br>$p = 0.096$ |
| Disease Duration (years) | | - | 10.97<br>(7.54) | 7.06<br>(5.07) | $F_{1,48} = 3.53$<br>$p = 0.066$ |
| Usual Pain Intensity (0-100) | | - | 53.91<br>(36.53) | 62.24<br>(32.86) | $F_{1,48} = 0.62$<br>$p = 0.434$ |
| Pain Catastrophizing Scale | | - | 26.59<br>(12.59) | 24.71<br>(11.42) | $F_{1,49} = 0.27$<br>$p = 0.606$ |
| Pain Anxiety Symptom Scale | | - | 92.09<br>(33.46) | 83.41<br>(30.42) | $F_{1,49} = 0.81$<br>$p = 0.373$ |
| Beck Depression Inventory-II | | - | 11.33<br>(8.29) | 19.12<br>(11.99) | $F_{1,47} = 7.06$<br>**$p = 0.011^*$** |

**Note:** Differences in categorical variables among groups were tested using the $\chi^2$ ; whereas differences in continuous variables were tested using ANOVA (Welch's ANOVA yielded very similar results).

**Table 2.**
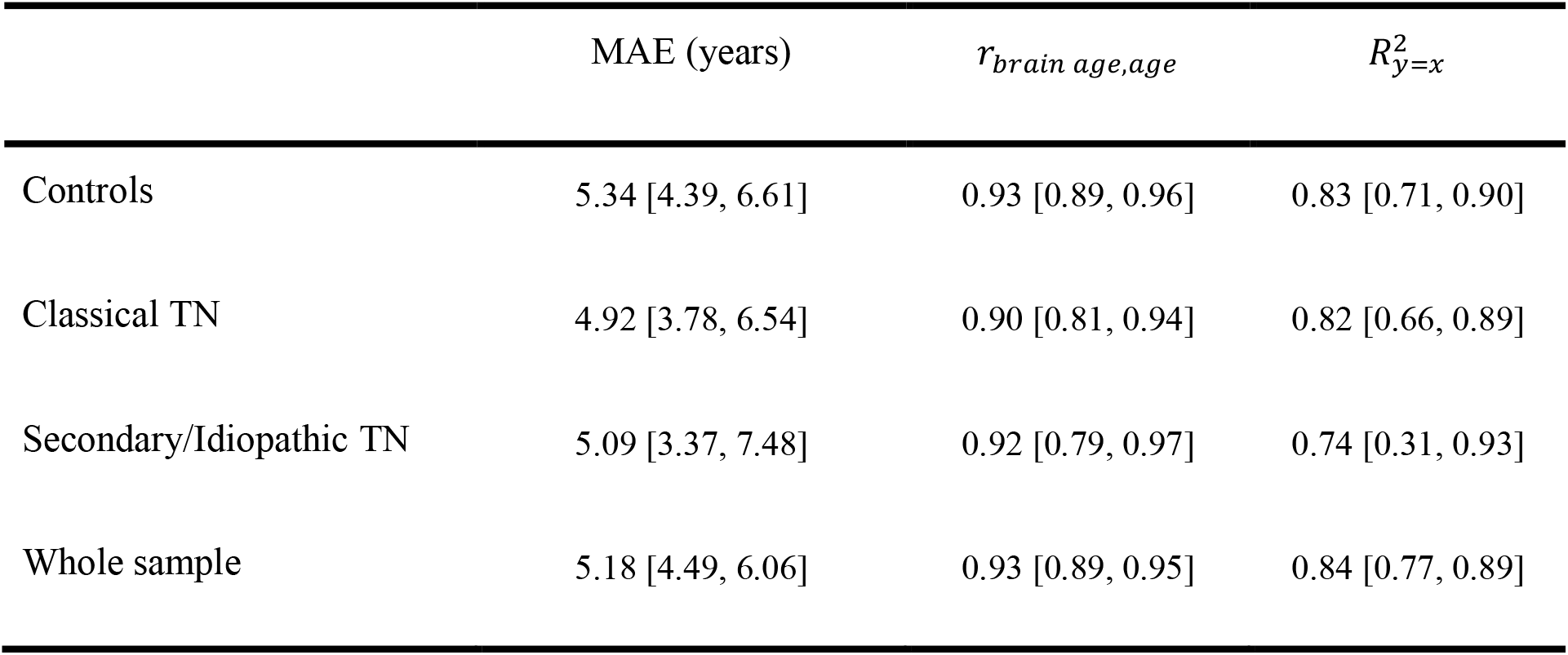
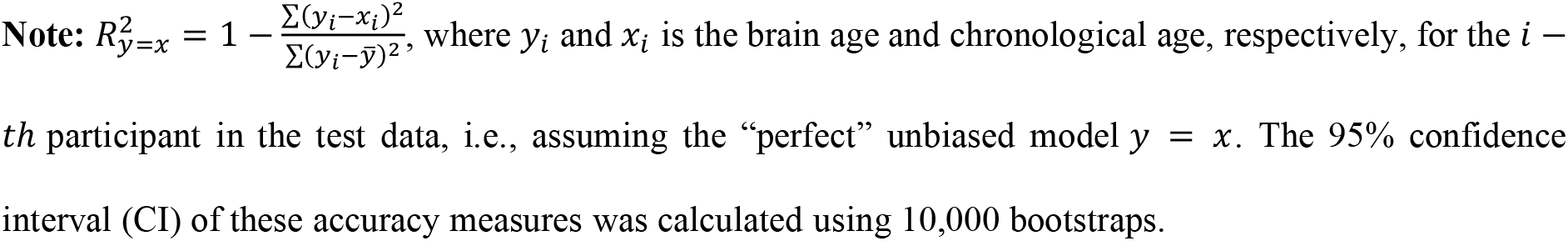
Accuracy of brain age predictions for each group and the whole sample.

| | MAE (years) | $r_{brain\ age, age}$ | $R^2_{y=x}$ |
| --- | --- | --- | --- |
| Controls | 5.34 [4.39, 6.61] | 0.93 [0.89, 0.96] | 0.83 [0.71, 0.90] |
| Classical TN | 4.92 [3.78, 6.54] | 0.90 [0.81, 0.94] | 0.82 [0.66, 0.89] |
| Secondary/Idiopathic TN | 5.09 [3.37, 7.48] | 0.92 [0.79, 0.97] | 0.74 [0.31, 0.93] |
| Whole sample | 5.18 [4.49, 6.06] | 0.93 [0.89, 0.95] | 0.84 [0.77, 0.89] |

**Note:** $R_{y=x}^2 = 1 - \frac{\sum(y_i - x_i)^2}{\sum(y_i - \bar{y})^2}$ , where $y_i$ and $x_i$ is the brain age and chronological age, respectively, for the $i$ -$th$ participant in the test data, i.e., assuming the “perfect” unbiased model $y = x$ . The 95% confidence interval (CI) of these accuracy measures was calculated using 10,000 bootstraps.

**Figure 1.**
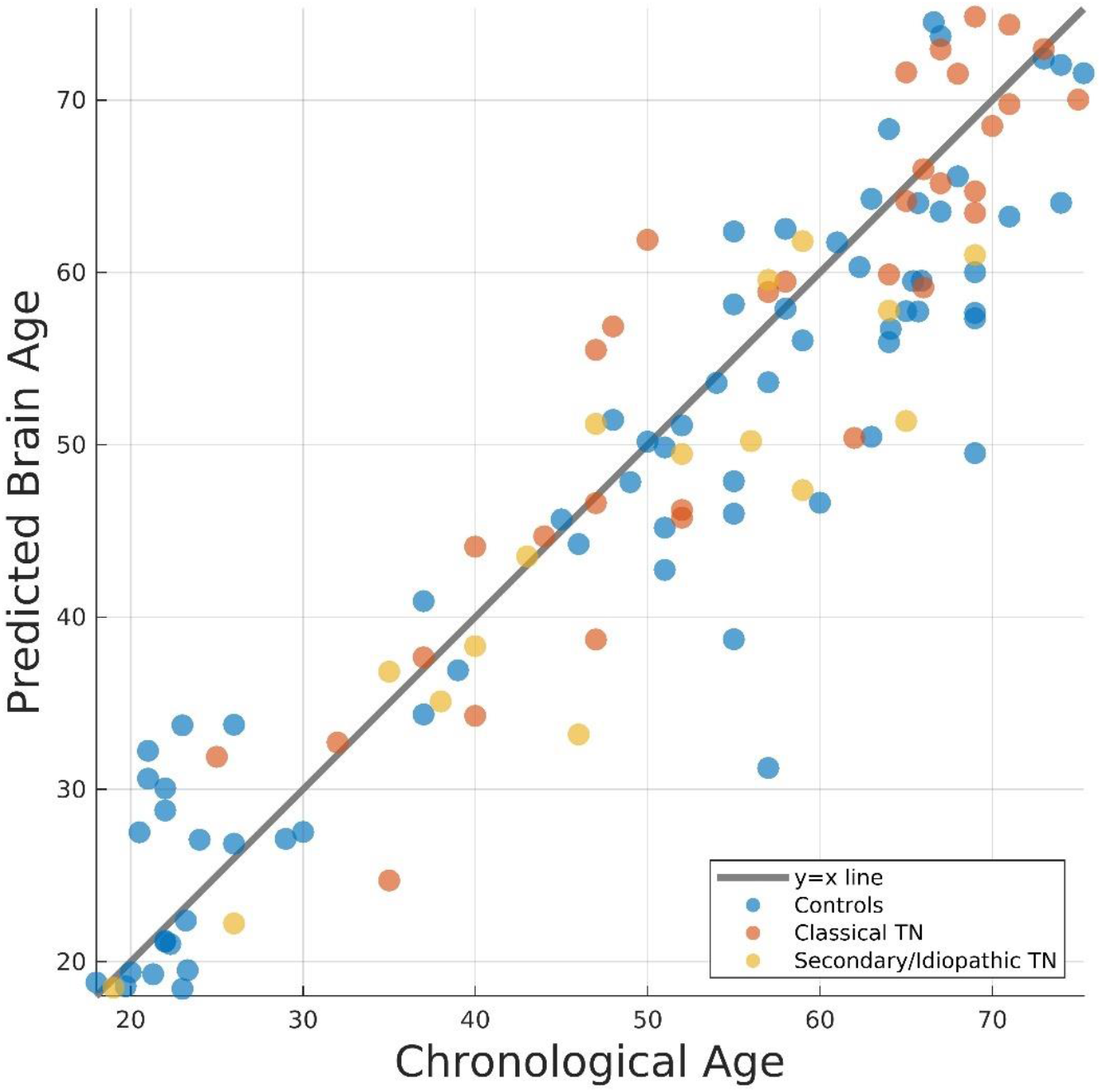
Brain age predictions versus chronological age for each group.

### 3.2. Differences in brain age acceleration between groups

To be consistent with the study by Hung and colleagues (2022), we first tested differences in brain-PAD between the TN and control groups. Brain-PAD was 2.5 years greater in TN patients compared to controls (p = 0.037, effect size f^2^ = 0.2^2^). Specifically, there were significant differences in brain-PAD among TN subtypes (ANCOVA p = 0.0078, effect size f^2^ = 0.28^2^). Indeed, for Classical TN, brain-PAD was significantly greater than the controls by 3.87 years (p = 0.01, Bonferroni-corrected across pairwise comparisons, **Figure 2**); whereas we found no evidence of a significant difference in brain-PAD between secondary/idiopathic TN and controls. When accounting for outliers, the results remained similar. Moreover, rejection of composite normality of the residuals failed in all fits (p>0.05). The group by sex interaction was not statistically significant (p = 0.664).

**Figure 2.**
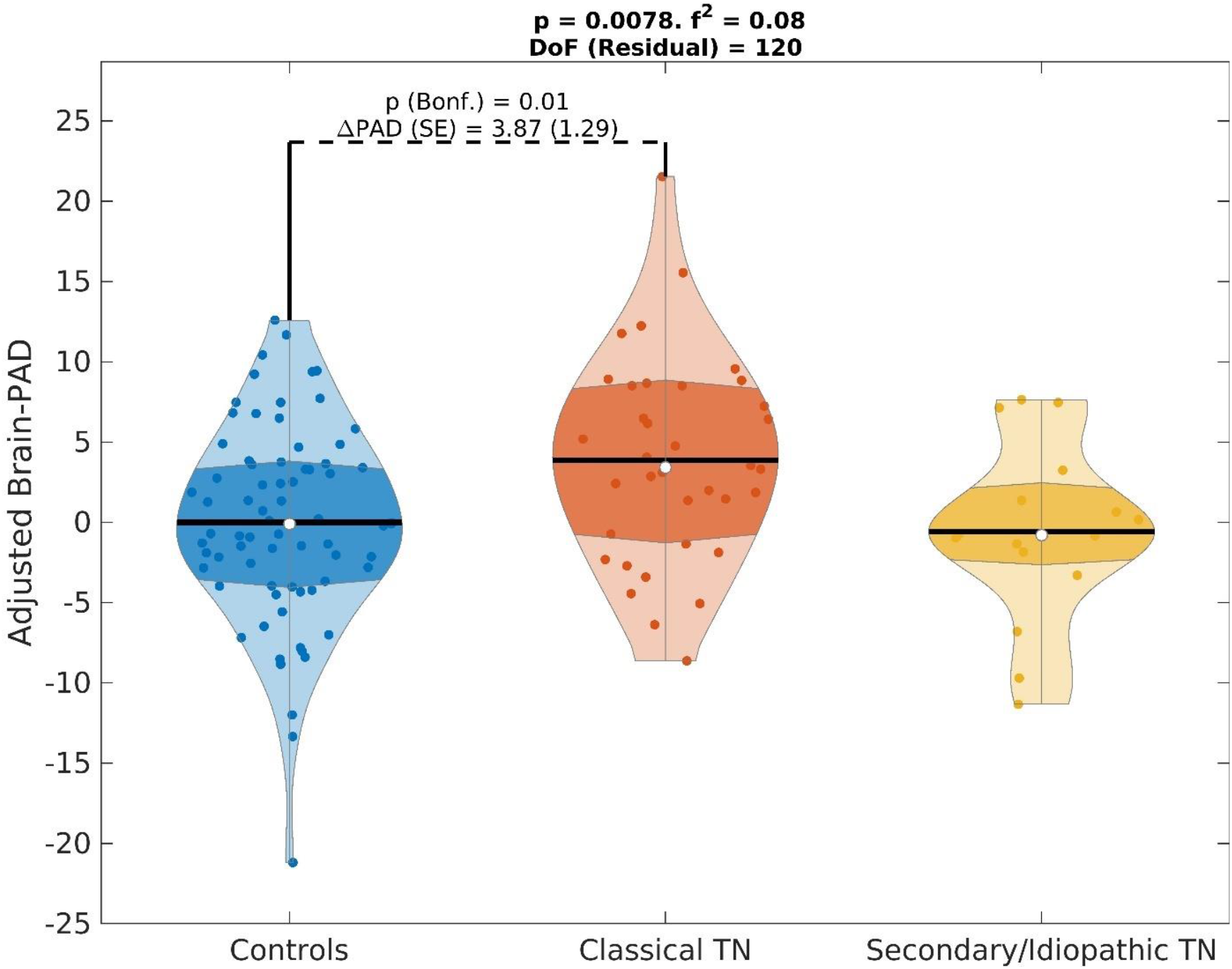
Differences in brain-PAD between groups, adjusting for age and sex. Within the violin plots, the shaded area is the interquartile region, the white dot indicates the median and the black horizontal line is the mean. Effect sizes, as quantified by Cohen’s f2, are also shown. DoF: Degrees of Freedom. ΔPAD (in years): difference in brain-PAD across groups. SE (in years): Standard Error.

The average (SE) of the unbiased brain-PAD (unbiased brain age minus chronological age) was −0.93 (0.82) years, 2.98 (1.21) years and −2.48 (1.46) years for the controls, classical TN, and secondary/idiopathic TN groups, respectively. Using a one-sample t-test, we found evidence that the average unbiased brain-PAD was significantly different from zero for the classical TN group (t_35_ = 2.47 with p = 0.019), whereas we did not found evidence for the other two groups (t_72_ = −1.12 with p = 0.26 and t_15_ = −1.70 with p = 0.11 for the controls and secondary/idiopathic TN, respectively). This suggests that the difference in brain-PAD is mainly driven by an accelerated brain age in the classical TN group as seen in **Figure 2**.

### 3.3. Association between brain age with clinical and psychological variables

Given that individuals with classical TN had significantly lower BDI-II scores, we ran partial correlations controlling for chronological age, sex, and BDI-II scores given the statistically significant differences in BDI-II scores between the TN subgroups. Brain-PAD had a significant positive association with both PCS (p = 0.032, corrected p = 0.041, effect size f^2^ = 0.33^2^) and PASS (p = 0.041, corrected p = 0.041, effect size f^2^ = 0.32^2^, **Figure 3**), but no significant association with disease duration (p = 0.519) or usual pain intensity (p = 0.443).

**Figure 3.**
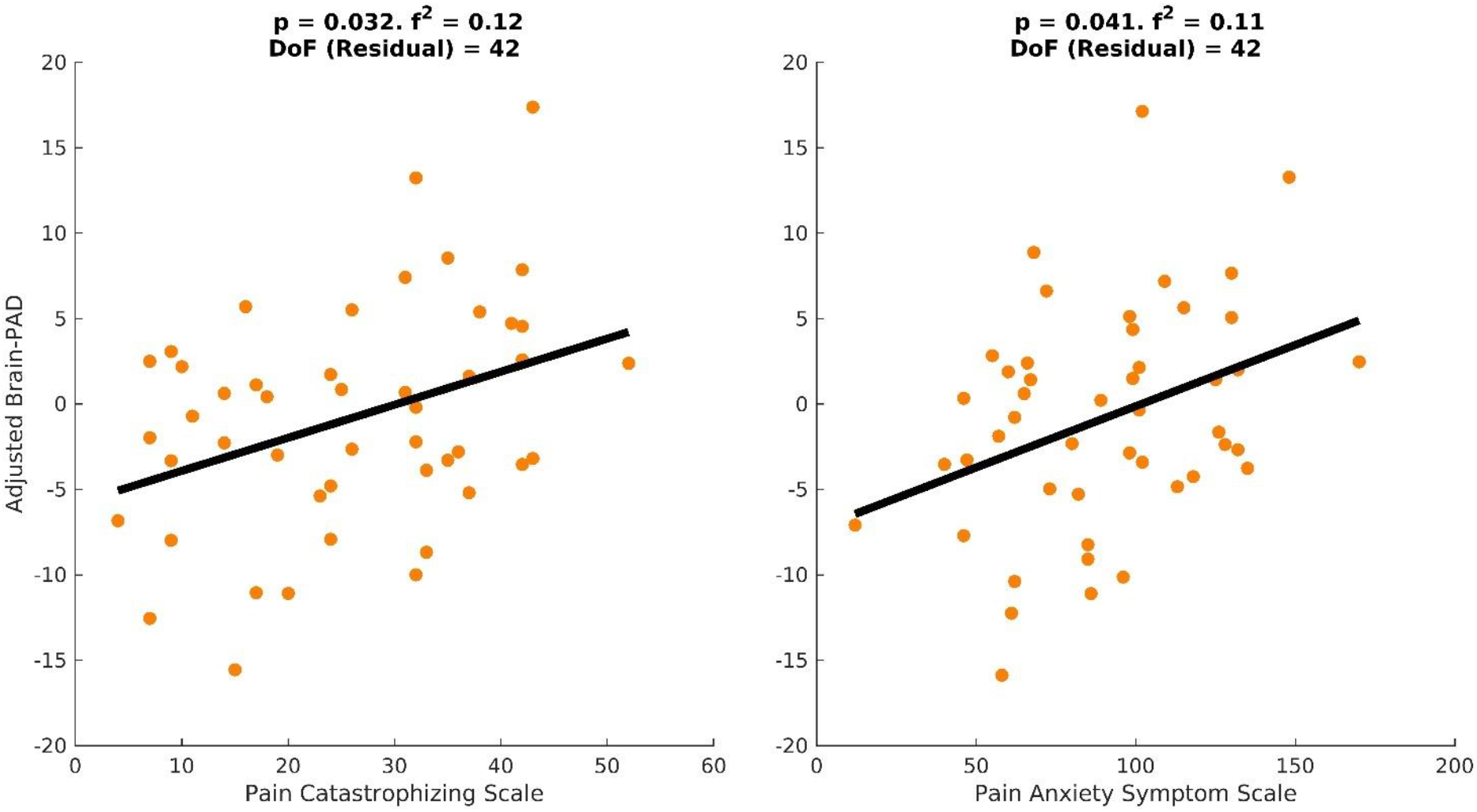
Scatter plots of brain-PAD versus scores of the Pain Catastrophizing Scale and the Pain Anxiety Symptom Scale.

## 4. Discussion

To our knowledge, this is the first study to examine a brain aging biomarker in persons with trigeminal neuralgia with consideration for the distinct types of trigeminal neuralgia diagnoses. Several important contributions emerged from this investigation. First, there were significant differences in brain-predicted age differences (brain-PAD or brain age gap) between individuals diagnosed with TN compared to age- and sex-matched pain-free controls. Second, those diagnosed with classical TN had “older” appearing brains compared to individuals with secondary/idiopathic TN or pain-free controls, regardless of age and sex. Finally, “older” brains were significantly associated with greater pain catastrophizing and pain-related anxiety levels in persons with TN, even after accounting for age, sex, and depressive symptoms.

Consistent with our hypothesis, individuals diagnosed with TN had 2.5 years “older” brain relative to their own individual’s chronological age adjusting for important covariates. This is consistent with the only study to date that has examined brain aging processes in persons with TN (Hung et al., 2022). However, that study did not differentiate between TN diagnoses. In our current study, it appears that brain aging was mainly driven by individuals diagnosed with classical TN with almost 4 years “older” brains relative to their own individual’s chronological age on average. Previous data suggests that each extra year of brain-predicted age (i.e., having a brain-PAD score of +1) results in a 6.1% relative subsequent increased mortality risk (Cole et al. 2018). Our findings also suggest that individuals diagnosed with secondary or idiopathic TN do not show accelerated brain aging processes as a group measured by our brain-wide brain aging biomarker. It is possible that in the classical TN presentation, the damage to the trigeminal nerve leads to brain aging processes either neuroinflammatory, neurodegenerative or both that are not a hallmark of secondary or idiopathic TN. This is currently speculative, and future studies both in rodents and humans are needed to elucidate the underlying brain aging mechanisms across the different TN subtypes as well as to test whether brain aging processes maybe antecedent or the result of the nerve compression.

An older brain age was also associated with greater pain catastrophizing and pain-related anxiety levels. This is consistent with our own previous work in older adults with musculoskeletal pain (Cruz-Almeida et al. 2019; Johnson et al. 2022). Interestingly, this remained statistically significant after accounting for differences in depressive symptoms, which is often co-morbid with pain, and anxiety across many chronic pain conditions. Taken together, our findings further support the importance of psychological characteristics in relation to brain structure and function that may be negatively impacted by chronic pain and be sensitive to our brain aging biomarker. Given that anxiety and depression have also been associated with accelerated brain aging (Johnson et al. 2022; Valdes-Hernandez et al. 2024) future research should attempt to disentangle these complex associations in persons with different TN diagnoses.

### 4.1 Limitations and future directions

Our study has some limitations. While the sample size for the training set was large, the TN study cohort was relatively small. However, TN participants are well-characterized in their TN characteristics including a certified clinician examination. Second, the current analysis was cross-sectional; therefore, we cannot determine whether a specific brain-predicted age preceded or was subsequent to TN-associated pain. From the present findings, directionality or causality cannot be inferred as it is equally possible that brain aging plays a central role for the sensitivity and resilience to many symptoms and disorders associated with biological aging processes including TN. Future longitudinal studies are needed to determine trajectories of brain aging and how they relate to TN and future health outcomes. Finally, our brain aging biomarker does not provide the anatomical specificity to determine which brain regions are specifically “aged” as brain aging processes are not a uniform process. Future studies are needed to further elucidate these associations. In addition, the development of region-specific aging biomarkers will help the field and ultimately clinical practice.

### 4.2 Conclusions

We report here accelerated brain aging processes in patients with classical TN, but not in persons diagnosed with secondary or idiopathic TN. Our study replicates previous findings in this population but adds to the literature that accelerated brain aging may not occur across all subtypes. Given the increased use of MRI for TN diagnostics, combined with our own recent work deriving our brain aging biomarker from clinical-grade scans, future studies within clinical settings are feasible and needed to truly understand this debilitating complex condition.

## Author contributions

Cruz-Almeida Y: Contributed to conception and design, contributed to analysis and interpretation, drafted the manuscript, critically revised the manuscript, gave final approval, agrees to be accountable for all aspects of work ensuring integrity and accuracy.

Valdes Hernandez P: Contributed to analysis and interpretation, drafted the manuscript, critically revised the manuscript, gave final approval, agrees to be accountable for all aspects of work ensuring integrity and accuracy.

Liang Y: Contributed to design, critically revised the manuscript, gave final approval, agrees to be accountable for all aspects of work ensuring integrity and accuracy.

Ding M: Contributed to design, contributed to interpretation, critically revised the manuscript, gave final approval, agrees to be accountable for all aspects of work ensuring integrity and accuracy.

Neubert JK: Contributed to design, contributed to interpretation, critically revised the manuscript, gave final approval, agrees to be accountable for all aspects of work ensuring integrity and accuracy.

All authors gave their final approval and agreed to be accountable for all aspects of the work.

## Acknowledgments

This work was supported by the Facial Pain Research Foundation (JN).

## References

Benjamini Y, Hochberg Y. 1995. Controlling the False Discovery Rate: A Practical and Powerful Approach to Multiple Testing. Journal of the Royal Statistical Society Series B (Methodological). 57(1):289–300. http://www.jstor.org/stable/2346101.

Cole JH, Ritchie SJ, Bastin ME, Valdés Hernández MC, Muñoz Maniega S, Royle N, Corley J, Pattie A, Harris SE, Zhang Q, et al. 2018. Brain age predicts mortality. Mol Psychiatry. 23(5):1385– 1392. [accessed 2018 Nov 16]. http://www.nature.com/doifinder/10.1038/mp.2017.62.

Cruz-Almeida Y, Fillingim RB, Riley JL, Woods AJ, Porges E, Cohen R, Cole JH, Riley III J, Adam W, Porges E, et al. 2019. Chronic Pain is Associated with a Brain Aging Biomarker in Community-Dwelling Older Adults. Pain. 160(5):1119–1130. [accessed 2019 Apr 28]. http://www.ncbi.nlm.nih.gov/pubmed/31009418.

Downton F, Blom G. 1961. Statistical Estimates and Transformed Beta-Variables. The Mathematical Gazette. 45(354):369.

Dozois DJA, Dobson KS, Ahnberg JL. 1998. A psychometric evaluation of the Beck Depression Inventory-II. Psychol Assess. 10(2):83–89.

Hung PSP, Zhang JY, Noorani A, Walker MR, Huang M, Zhang JW, Laperriere N, Rudzicz F, Hodaie M. 2022. Differential expression of a brain aging biomarker across discrete chronic pain disorders. Pain. 163(8):1468–1478. [accessed 2024 May 27]. https://pubmed.ncbi.nlm.nih.gov/35202044/.

Johnson AJ, Buchanan T, Laffitte C, Huo Z, Cole JH, Buford TW, Fillingim RB, Cruz-Almeida Y. 2022. High impact chronic knee pain is associated with greater clinical and experimental pain and predicted brain aging in middle-aged and older adults. Exp Gerontol.(Under Review).

McCracken LM, Zayfert C, Gross RT. 1992. The Pain Anxiety Symptoms Scale: development and validation of a scale to measure fear of pain. Pain. 50(1):67–73. [accessed 2024 May 27]. https://pubmed.ncbi.nlm.nih.gov/1513605/.

Neter J. 1990. Applied linear statistical models. Regression, analysis of variance, and statistical designs.

Olesen J. 2018. Headache Classification Committee of the International Headache Society (IHS) The International Classification of Headache Disorders, 3rd edition. Cephalalgia. 38(1):1–211. [accessed 2024 May 27]. https://pubmed.ncbi.nlm.nih.gov/29368949/.

Shapiro SS, Wilk MB. 1965. An analysis of variance test for normality (complete samples). Biometrika. 52(3–4):591–611.

Sullivan MJL, Bishop SR, Pivik J. 1995. The Pain Catastrophizing Scale: Development and Validation. Psychol Assess. 7(4):524–532.

Valdes-Hernandez PA, Johnson AJ, Montesino-Goicolea S, Laffitte Nodarse C, Bashyam V, Davatzikos C, Fillingim RB, Cruz-Almeida Y. 2024. Accelerated Brain Aging Mediates the Association Between Psychological Profiles and Clinical Pain in Knee Osteoarthritis. J Pain. 25(5). [accessed 2024 May 27]. https://pubmed.ncbi.nlm.nih.gov/37952863/.

Valdes-Hernandez PA, Laffitte Nodarse C, Cole JH, Cruz-Almeida Y. 2023. Feasibility of brain age predictions from clinical T1-weighted MRIs. Brain Res Bull. 205. [accessed 2024 May 27]. https://pubmed.ncbi.nlm.nih.gov/37952679/.

Valdes-Hernandez PA, Laffitte Nodarse C, Johnson AJ, Montesino-Goicolea S, Bashyam V, Davatzikos C, Peraza JA, Cole JH, Huo Z, Fillingim RB, et al. 2023. Brain-predicted age difference estimated using DeepBrainNet is significantly associated with pain and function-a multi-institutional and multiscanner study. Pain. 164(12):2822–2838. [accessed 2024 May 27]. https://pubmed.ncbi.nlm.nih.gov/37490099/.

Valdes-Hernandez PA, Laffitte Nodarse C, Peraza JA, Cole JH, Cruz-Almeida Y. 2023. Toward MR protocol-agnostic, unbiased brain age predicted from clinical-grade MRIs. Sci Rep. 13(1). [accessed 2024 May 27]. https://pubmed.ncbi.nlm.nih.gov/37950024/.

Zhang C, Hu H, Das S, Yang M-J, Li B, Li Y, Xu X, Yang H-F. 2020. Structural and Functional Brain Abnormalities in Trigeminal Neuralgia: A Systematic Review. J Oral Facial Pain Headache. 34(3):222–235.

